# Chitosan perception in *Arabidopsis* requires the chitin receptor AtCERK1 suggesting an improved model for receptor structure and function

**DOI:** 10.1101/170092

**Authors:** Ekaterina Gubaeva, Airat Gubaev, Rebecca Melcher, Stefan Cord-Landwehr, Ratna Singh, Nour Eddine El Gueddari, Bruno M. Moerschbacher

## Abstract

Chitin, a linear polymer of N-acetyl-D-glucosamine, and chitosans, fully or partially deacetylated derivatives of chitin, are known to elicit defense reactions in higher plants. We compared the ability of chitin and chitosan oligomers and polymers (chitin oligomers with degree of polymerization 3 to 8; chitosan oligomers with degree of acetylation 0% to 35% and degree of polymerization 3 to 15; chitosan polymers with degree of acetylation 1% to 60% and degree of polymerization ~1300) to elicit an oxidative burst indicative of induced defense reactions in *A. thaliana* seedlings. Fully deacetylated chitosans were not able to trigger a response; elicitor activity increased with increasing degree of acetylation of chitosan polymers. Partially acetylated chitosan oligomers required a minimum degree of polymerization of 6 and at least four N-acetyl groups to trigger a response. Invariably, elicitation of an oxidative burst required the presence of the chitin receptor AtCERK1. Our results as well as previously published studies on chitin and chitosan perception in plants are best explained by a new general model of LysM-containing receptor complexes where two partners form a long, but off-set chitin-binding groove and are, thus, dimerized by one chitin or chitosan molecule, sharing a central GlcNAc unit with which both LysM domains interact. To verify this model and to distinguish it from earlier models, we assayed elicitor and inhibitor activities of selected partially acetylated chitosan oligomers with fully defined structures. In contrast to the initial “continuous groove”, the original “sandwich”, or the current “sliding mode” models for the chitin/chitosan receptor, the here proposed “slipped sandwich” model - which builds on these earlier models and represents a consensus combination of these - is in agreement with all experimental observations.

## Introduction

Chitin is an evolutionary ancient, phylogenetically widespread biopolymer present throughout most realms of eukaryotes, with the notable exception of higher plants and vertebrates in both of which it, consequently, acts as a convenient trigger for immune responses upon non-self recognition. While it is widely accepted that chitin receptors in plants are multicomponent transmembrane complexes, the components of which are related but differ in detail between species, and while crystal structures have been solved and models have been built, a number of questions are still debated controversially. Which components make up the receptors? How do these components bind the chitin ligands? How does chitin binding lead to receptor complex formation?

Chitin perception is best studied in *Arabidopsis* and rice (*Oryza sativa*), where similar but not identical chitin receptor complexes have been described (Figure 1, ^1–4^, reviewed in ^5,6^). AtCERK1, a transmembrane protein with three extracellular chitin-binding LysM domains and an intracellular protein kinase domain, is known as the central component of the chitin receptor in *Arabidopsis* (Figure 1A, ^1^), possibly in conjunction with AtLYK5 and/or AtLYK4 (Figure 1B, ^2^). In contrast, OsCERK1 in *O. sativa* does not bind chitin but rather interacts with OsCEBiP that recognizes chitin, a related protein with three LysM domains but lacking the protein kinase domain (Figure 1C ^3,4,7,8^). In both *Arabidopsis* and rice, interaction with chitin octamer appears to lead to dimerization and activation of the receptor. Two alternative hypotheses have been put forward in *Arabidopsis*. Either binding of the chitin octamer directly dimerizes AtCERK1 (Figure 1A), ^1^ or it may lead to the formation of a hetero-tetramer of a preformed AtLYK5 dimer and two AtCERK1 moieties (Figure 1B). ^2^ In rice, a preformed hetero-dimer of OsCERK1 and OsCEBiP is believed to dimerize upon chitin octamer binding, which brings together also two OsCERK1 proteins, resulting in downstream signaling (Figure 1C). ^3,4^

**Figure 1.**
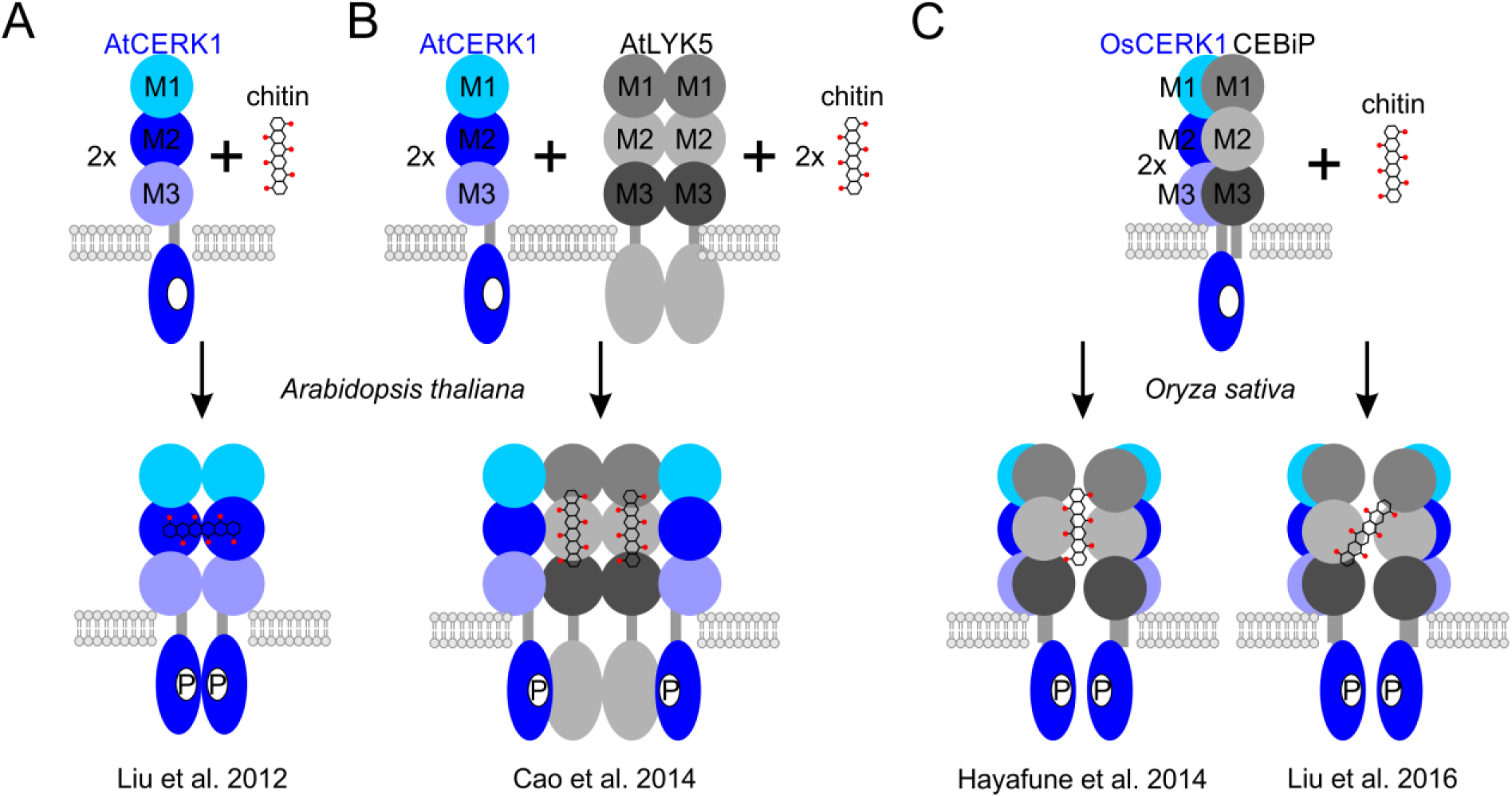
Models of chitin receptors and their activation. **A**: Chitin-induced AtCERK1 dimerization in *Arabidopsis*. Direct binding of a chitin oligomer to AtCERK1’s LysM2 domain leads to homo-dimer formation and AtCERK1 activation. **B**: Chitin-induced hetero-tetramer receptor complex in *Arabidopsis*. Two AtLyK5 forming a homo-dimer bind two chitin oligomers by all three LysM domains and recruit two AtCERK1 to build the active receptor complex. **C**: Chitin-induced hetero-tetramer receptor complex in rice. OsCERK1 and OsCEBiP form a hetero-dimer; a chitin oligomer either “sandwiches” between two OsCEBiP proteins leading to the formation of the active receptor complex or, alternatively, the chitin oligomer induces homo-dimerization of two OsCEBiP molecules in the “sliding mode”. In both cases, elevated levels of association of OsCEBiP with OsCERK1 in the presence of chitin lead to OsCERK1 dimerization and activation.

There is also controversy concerning how the receptor binds chitin. The crystal structure of the LysM domains of AtCERK1 revealed four ligand binding subsites on subunit LysM2 (Figure E1, ^1^), three of them directly interacting with GlcNAc residues. Recently, the crystal structure (Figure E1, ^4^) of OsCEBiP revealed that chitin is bound in a similar way as seen in the AtCERK1-chitin complex. In both cases, the chitin oligomer appears to bind to LysM2 only, the other two LysM domains are not involved in ligand interaction. An alternative mode of chitin binding has been shown for Ecp6, the LysM containing effector of the fungal tomato pathogen *Cladosporium fulvum*, which binds chitin by intramolecular LysM dimerization, involving LysM1 and LysM3 (Figure E1, ^9^). Interestingly, the model build for chitin binding to AtLYK4/AtLYK5 in *Arabidopsis* was based on the Ecp6 model, suggesting a different conformation and a different mode of ligand interaction compared to AtCERK1 ^2^.

**Figure E1.**
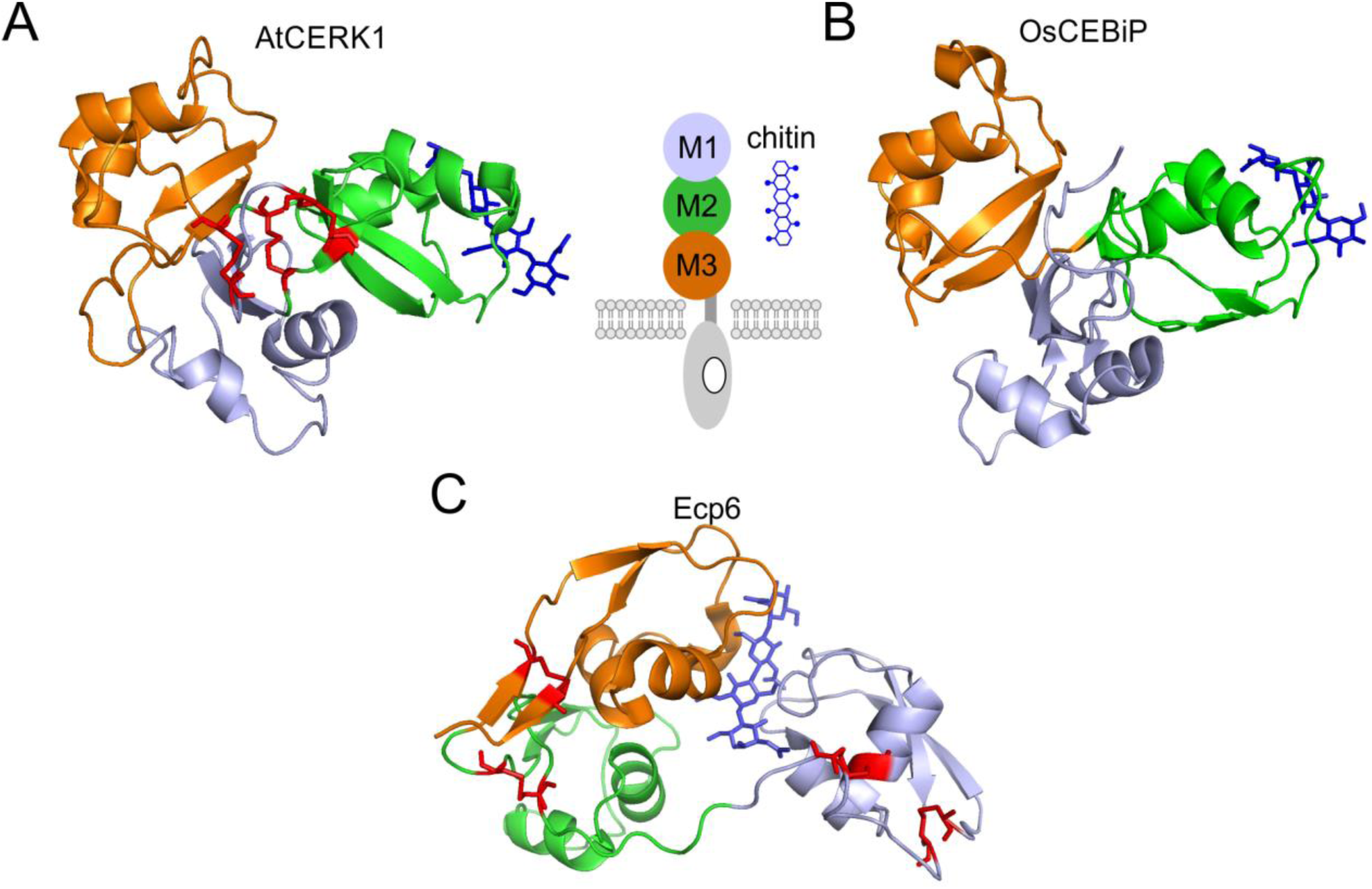
Crystal structures of AtCERK1 (PDB ID 4EBZ) (**A**) and OsCEBiP (PDB ID 5JCE) (**B**) ectodomains as well as of the fungal chitin-binding protein Ecp6 (PDB ID 4B8V) (**C**). The structures of the AtCERK1 and OsCEBiP ectodomains are very similar and in both cases, chitin is bound to the surface of the LysM2 domain. In contrast, Ecp6 assumes a different conformation, allowing for chitin binding by intra-chain LysM (M1 and M3) dimerization deeply buried inside the protein. Disulfide bonds in Ecp6 and AtCERK1 are indicated in red. LysM domains are colored as LysM1: light blue; LysM2: green; LysM3: orange. The chitin molecule is indicated by blue colored sticks.

Similarly, different models have been proposed for the mechanism of chitin-induced receptor dimerization. The observation of four GlcNAc binding sites in AtCERK1 suggested a simple, “continuous groove” model in which a chitin octamer, which is considered the most elicitor-active chitin oligomer, would span and bridge two receptor molecules, dimerizing them ^1^. In rice, however, based on sophisticated STD NMR studies, it was first suggested that chitin is “sandwiched” between the two OsCEBiP receptor units, connecting them via the acetyl groups oriented alternatingly to both sides (Figure 1C), ^3^ but solving of the crystal structure (Figure E1) ^4^ led to the “sliding mode” model for receptor dimerization, based on an alignment to the LysM domain in the chitin hexamer-mediated NlpC/P60-LysM homo-dimer formation of an endopeptidase ^10^, where three GlcNAc residues are bound by the first OsCEBiP molecule and the next three GlcNAc residues are bound by the second. This “sliding mode” model, thus, is a combination of the earlier “continuous groove” and “sandwich” models.

Clearly, the AtCERK1 and OsCEBiP binding sites accommodate only four GlcNAc residues of which three interact directly with the LysM domain ^1,4^ which seems at odds with the necessity of a chitin octamer for optimal elicitor activity ^1,8,11,12^ if the “sandwich” model ^3^ applies. Similarly, the “sliding mode” model ^4^ does not explain why chitin octamer is a much better elicitor than chitin hexamer ^18,11,12^. In contrast, the “continuous groove” model ^1^ does not explain why a chitosan octamer consisting of alternating GlcN and GlcNAc residues acts as an inhibitor of chitin octamer-induced elicitation rather than as an elicitor ^3^.

To address these questions, we tested a larger set of chitosan oligomers and polymers differing in their degree of polymerization (DP) and degree of acetylation (DA) for their elicitor activities, both in wild type *Arabidopsis* seedlings and in knock-out mutants lacking the AtCERK1 chitin receptor. Plants are known to recognize chitosans, i.e. partially de-*N*-acetylated chitin, but the mechanism of recognition is unknown ^13,14^. It has been speculated that electrostatic interactions may be involved ^15^ or plants may contain chitosan-specific receptors ^16,17^. Alternatively, it has been suggested that the chitin receptor may also recognize chitosan provided its DA is high enough ^18^. We here show that AtCERK1 is indeed required for the elicitation of an oxidative defense reaction in *Arabidopsis* by a broad range of chitosan polymers and oligomers, and based on these experimental results, we propose an improved model for chitin-and chitosan-induced dimerization of the chitin receptor in *Arabidopsis* which should also be applicable for chitin and chitosan perception in rice.

## Results and Discussion

Wild-type *A. thaliana* seedlings treated with chitin oligosaccharides clearly showed a size-(DP) and dose-dependent induction of H_2_O_2_ (Figure 2A), as expected ^1,11,12^. Starting from DP of 6, we observed an increase in ROS generation with increasing DP at the concentration of 1 μg/ml; the maximum of eliciting activity was observed for chitin oligomers of DP 7 and DP 8, confirming published results ^1,2^. Also as expected, the *Atcerk1* mutant seedlings did not show any response to chitin even at 10 μg/ml, thus corroborating that AtCERK1 is involved in recognition of chitin oligomers ^18–21^, and validating the assay. Fully deacetylated chitosan oligomers (DA 0%, DP 3-8) did not trigger an immune response in *A. thaliana* seedlings (Figure 2B).

**Figure 2.**
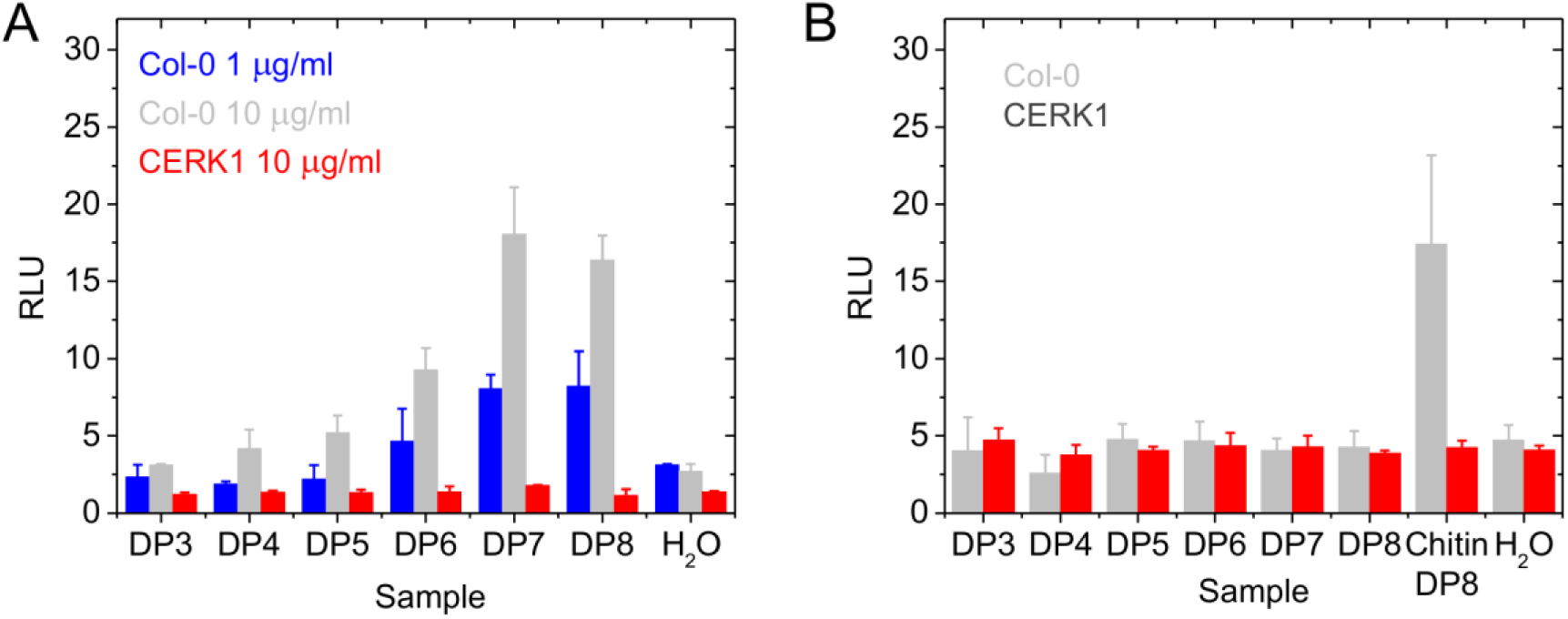
Validation of the assay method to quantify chitin-induced ROS production (H_2_O_2_) in *A. thaliana*. **A**. ROS production by wild-type (Col-0) and *Atcerk1* (CERK1) seedlings in response to the addition of chitin oligomers with different DP (from 3 to 8). **A**: Starting from DP 6, chitin oligomers induced ROS formation that was strongest for DP 7 and DP 8, at a concentrations as low as 1 μg/ml. ROS formation increased with increasing concentration of chitin oligomers. The *Atcerk1* mutant did not show any detectable ROS response caused by addition of chitin oligomers. **B**: Fully deacetylated chitosan oligomers with different DP (from 3 to 8) did not elicit ROS production in wild-type (Col-0) or *Atcerk1* (CERK1) seedlings; ROS formation in the presence of chitin oligomer with DP 8 served as positive control. Water addition to the seedlings served as negative control. Data given are means ± SD of four replicates from one, representative of three independent experiments.

All partially acetylated chitosan polymers tested, spanning a wide range of DA from 1 to 60%, elicited an oxidative burst; elicitor activity increased gradually with increasing DA (Figure 3A). In all cases, the elicitor activity of the chitosans was dependent on the presence of AtCERK1, arguing against disturbance of plant membranes through electrostatic interactions as a perception mechanism for chitosan polymers which had previously been proposed ^15^.

**Figure 3.**
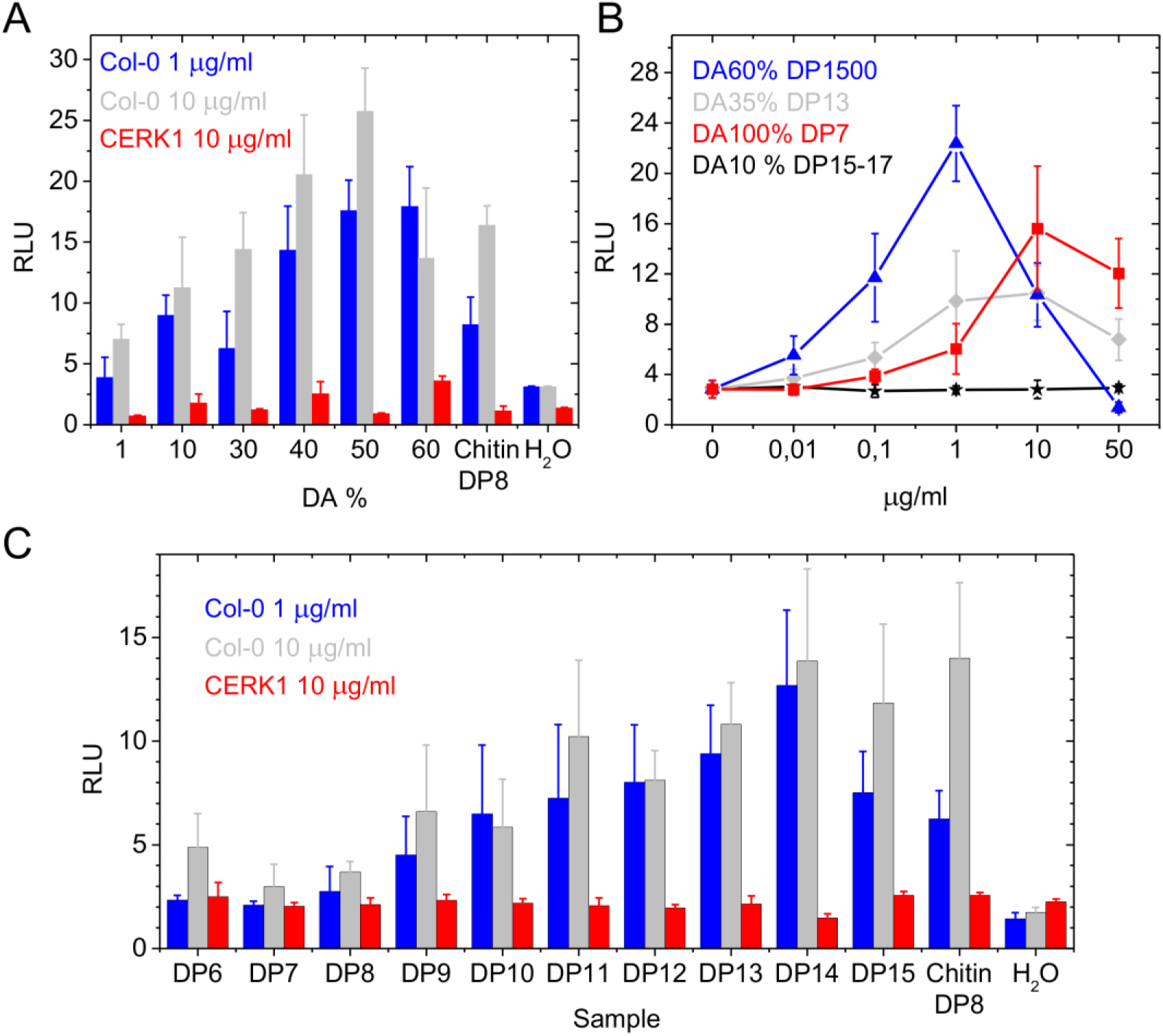
Elicitor activity of partially acetylated chitosan polymers and oligomers. **A**. ROS production by wild-type (Col-0) and *Atcerk1 A. thaliana* seedlings in response to the addition of chitosan polymers with varying DA (from 1% to 60%) and constant DP (around 1300). Elicitor activity increased with increasing DA and was similar to the positive control (chitin octamer) already at rather low DA. The *Atcerk1* mutant did not show any detectable ROS response caused by addition of chitosan in comparison to the water control. **B**. Dependence of ROS formation on the concentration of chitosan oligomers and polymers (from 0 (water control) to 50 μg/ml) with different DP and DA in wild-type *A. thaliana* seedlings. ROS formation increased with increasing chitosan concentration until it reached a maximum beyond which ROS formation decreased, except for very low DA chitosan oligomers which did not elicit any detectable ROS production. **C**. ROS production by wild-type (Col-0) and *Atcerk1 A. thaliana* seedlings in response to the addition of chitosan oligomers with different DP (from 6 to 15) and constant DA of 35%. Chitosan oligomers of DP 9 and larger induced AtCERK1-dependent ROS formation that increased with increasing DP. Data given are means ± SD of four replicates from one, representative of three independent experiments.

A minimum DP of 9 was required when chitosan oligomers of DA 35% were used as elicitors, and a maximum response was reached with DP 14 (Figure 3C). In contrast, chitosan oligomers of DA 10% were elicitor-inactive in the DP range investigated (from 4 to 15-17) (Figure E3). Again, the *Atcerk1* mutant seedlings did not show any response when treated with the chitosan oligomers, arguing against chitin-receptor-independent perception mechanisms for chitosans which also had previously been proposed ^15^. Dose-response curves showed that elicitor activity increased with increasing DA and increasing DP. At higher concentrations, elicitor activities tended to decrease, indicating either a toxic effect of the chitosans to the seedlings or a lack of ligand-induced receptor dimerization at super-optimal ligand concentrations. Interestingly, chitosan oligomers of DA 35% and DP 13 had slightly higher elicitor activities at low concentrations than fully acetylated chitin oligomers of DP 7 (Figure 3B), indicating that longer molecules may be more efficient in activating the receptor complex. Presumably, such molecules require a minimum number - and possibly distribution - of acetyl groups for efficient binding, and a minimum number of free amino groups for solubility.

**Figure E3.**
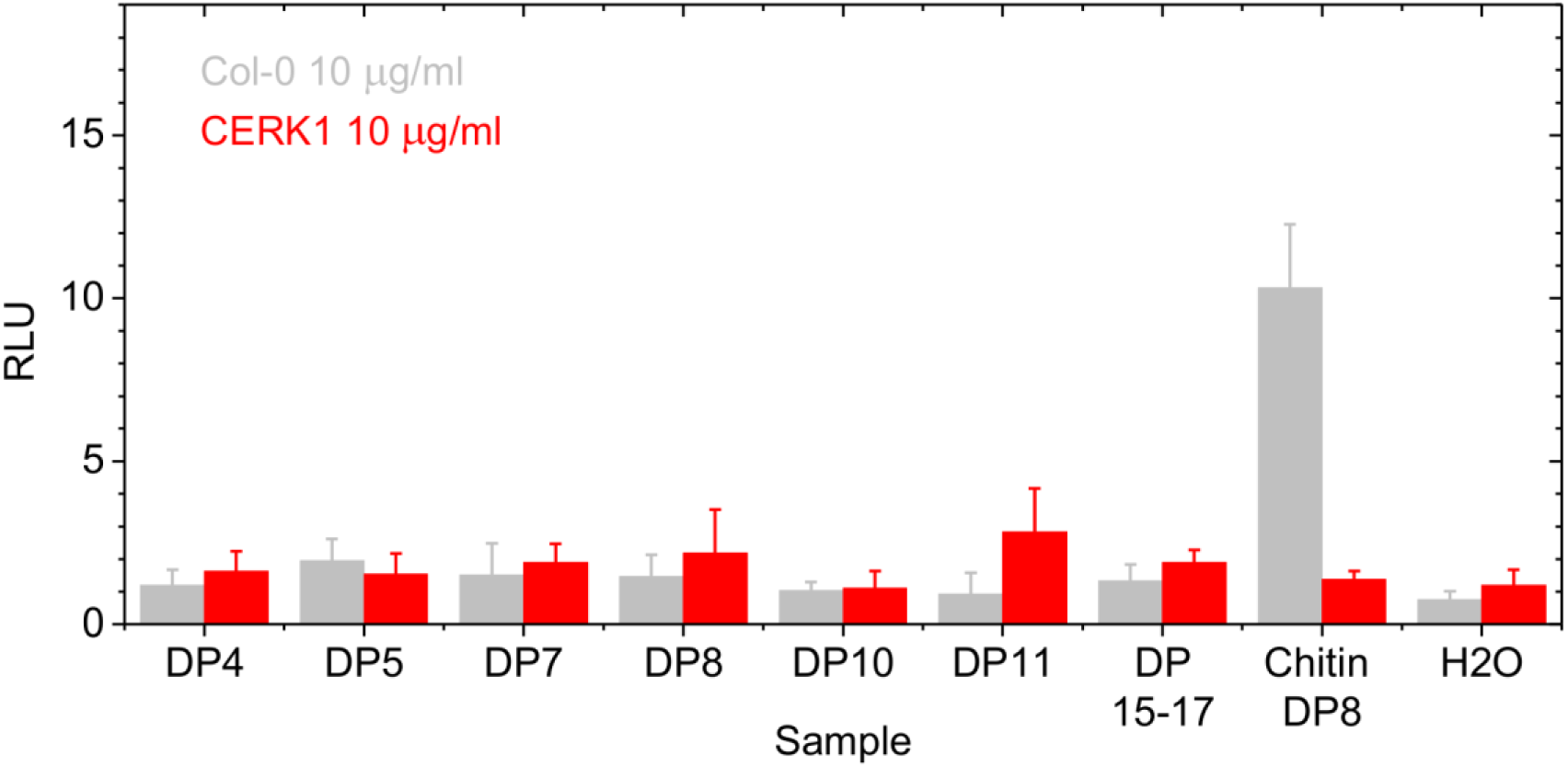
Elicitor activitiy of partially acetylated chitosan oligomers. ROS production by wild-type (Col-0) and *Atcerk1 A. thaliana* seedlings in response to the addition of chitosan oligomers with different DP (from 4 to 15-17) and constant DA of 10%. No ROS formation was induced by these oligomers. Chitin oligomer with DP 8 served as positive control. Addition of water instead of chitin served as negative control. Data given are means ± SD of four replicates from one, representative of three independent experiments.

Chitosan oligomers with DA 10% and DP 17, which on average contain less than two acetyl groups (17 × 10% = 1.7) per oligomer, did not induce ROS generation (Figure E3) although they fulfill the minimum length requirement for chitin oligomer perception (DP ≥ 6, Figure 2A). In contrast, chitosan oligomers with DA 35% and DP 9, which on average contain more than three (9 × 35% = 3.2) acetyl groups per molecule, did induce a ROS response (Figure 3C). Therefore, to induce ROS generation, we assume that a minimum of four acetyl groups would be required in a chitosan hexamer to show elicitor activity.

In an attempt to explain our experimental results on the basis of the models proposed for chitin perception in plants, we had a closer look at the crystal structure of AtCERK1 in complex with chitin pentamer ^1^. When we tried to model a dimer consisting of two AtCERK1 molecules with a chitin tetramer bound, we were surprised to find that this is possible without steric clashes between the two receptors if we superimpose the GlcNAc residue at the reducing end (“NAG-4”) of one tetramer and the GlcNAc residue at the non-reducing end (“NAG-1”) of the other tetramer (Figure 4). Thus, two chitin tetramers are aligned and form one chitin heptamer. This model for chitin-induced AtCERK1 dimerization which we termed the “slipped sandwich” model combines the original “sandwich” model ^3^ and the recently described “sliding mode” model ^4^ for chitin-induced OsCEBiP dimerization. It differs from the “sandwich model” by offsetting the two receptor molecules by three rather than just one glycosyl residue, leading to just one instead of three shared residues binding to both receptor molecules. It differs from the “sliding mode” model by taking into account four instead of three binding subsites per receptor molecule, leading to dual binding of the central GlcNAc unit which is not seen in the “sliding mode” model. In contrast to the “sliding mode” model, the “slipped sandwich” model explains why chitin heptamer is a much better elicitor than chitin hexamer. In contrast to the “continuous grove” model, it also explains why chitin heptamer and chitin octamer have equally strong elicitor activities. Of course, the situation may be different in *Arabidopsis* and rice, but the “slipped sandwich” model seems to be consistent with all experimental observations made in both systems and, thus, might apply to chitin perception in both species.

**Figure 4.**
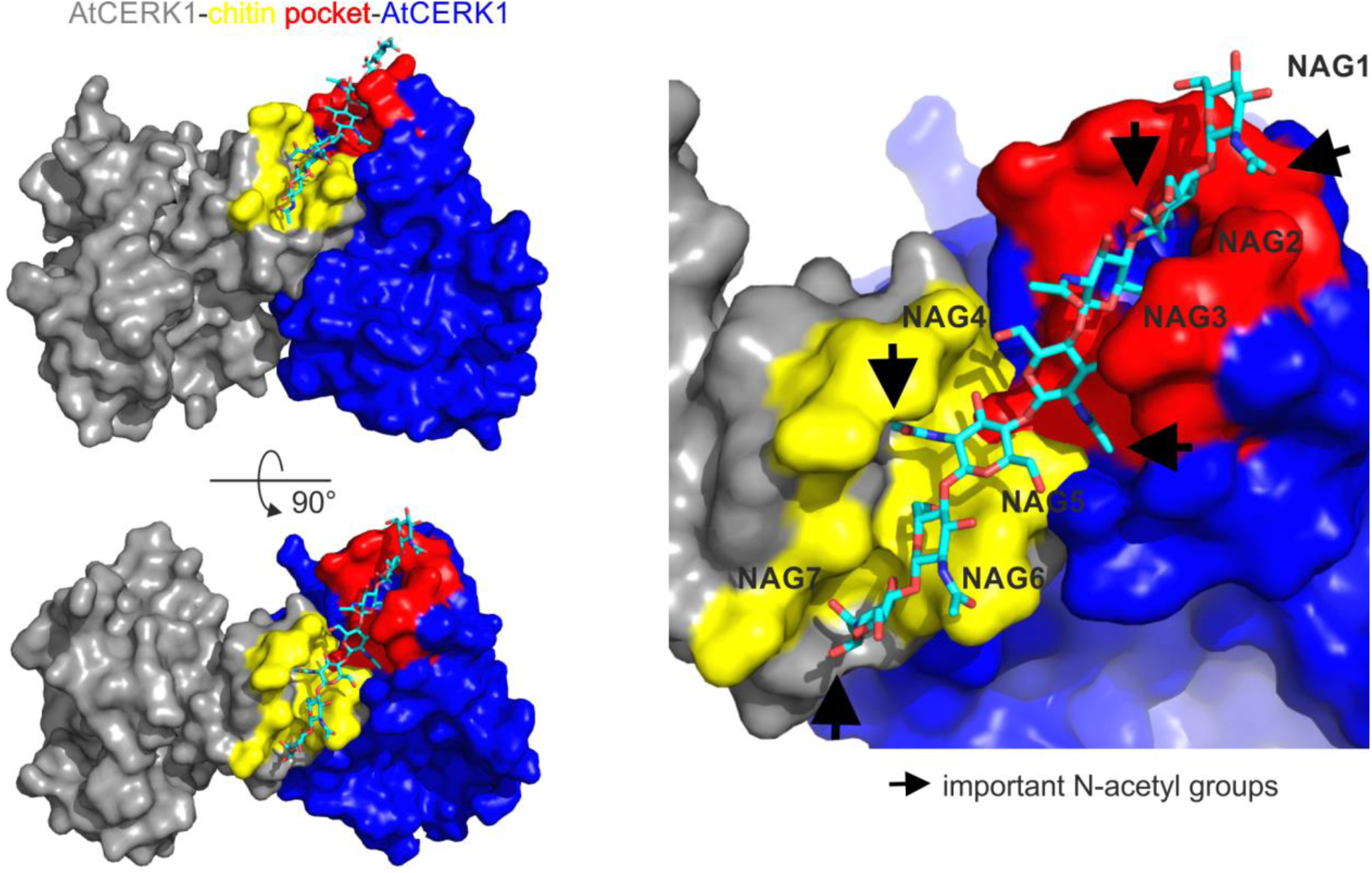
Model of chitin-induced AtCERK1-AtCERK1 ectodomain homo-dimerization. One AtCERK1 is depicted in gray (PDB ID 4EBZ), the second AtCERK1 in blue. The binding pocket is formed by the LysM2 domains of both AtCERK1 ectodomains. The part of the binding pocket formed by the gray AtCERK1 is depicted in yellow, that formed by the blue AtCERK1 in red. To build the chitin heptamer, shown in sticks representation, from two chitin tetramers, each positioned as seen in the crystal structure together with its corresponding AtCERK1 ectodomain ^1^, we aligned the non-reducing end GlcNAc residue of one of the tetramers to the same position (NAG4 in heptamer) as the reducing end GlcNAc unit of the second tetramer. Arrows indicate possible important N-acetyl groups for binding (NAG1, NAG2, NAG4, NAG5, NAG7).

The model would also allow partially acetylated chitosans and chitosan oligomers to bind and dimerize the receptor as long as their pattern of acetylation is such that enough acetyl groups are facing the binding site on the receptors while free amino groups are oriented away from the receptors. In fact, when analyzing the ligand-bound AtCERK1 and the OsCEBiP structures, we realized that of the four GlcNAc subunits interacting directly or indirectly with the LysM domains, the acetyl group of the third GlcNAc from the non-reducing end points away from the receptor and, thus, makes no interaction with it. Consequently, the mono-deacetylated chitosan tetramer GlcNAc-GlcNAc-GlcN-GlcNAc should bind equally well as the fully acetylated chitin tetramer. As a consequence, experiments using partially acetylated chitosan oligomers with fully defined structure in terms of DP, DA, and PA (pattern of acetylation) might help to critically assess the different models for ligand binding and receptor dimerization.

A chitosan oligomer consisting of alternating GlcN and GlcNAc units has been shown to act as an inhibitor of chitin perception in rice ^3^. In the original “sandwich” model ^3^, this was explained by the oligomer binding to one OsCEBiP molecule but then being unable to bind a second OsCEBiP molecule (Figure 5A). In the “continuous groove” model ^1^, two OsCEBIP molecules would align to form a continuous binding groove with 2 × 4 = 8 binding sites, and the alternating chitosan oligomer would be expected to be able to bind, as all acetyl groups would point towards one side of the oligomer, thus dimerizing the receptor - in which case this chitosan oligomer should act as an elicitor rather than as an inhibitor. The “continuous groove” model, thus, is not compatible with this experimental result, in contrast to the other three models. In case of the “sliding mode” model ^4^, the LysM2 domain of the first OsCEBiP molecule could bind to a motif within this oligomer, but the second OsCEBiP molecule could then not bind in the required close proximity to the first one to lead to dimerization. Hence, in this model, the alternating chitosan octamer would act as an inhibitor, as observed. The same would be true in our “slipped sandwich” model - which differs from the “sliding mode” model only in the presence of the shared middle GlcNAc unit of chitin as well as in four rather than three GlcNAc units bound per LysM domain, and from the original “sandwich” model in being staggered three instead of just one glycosyl unit.

**Figure 5.**
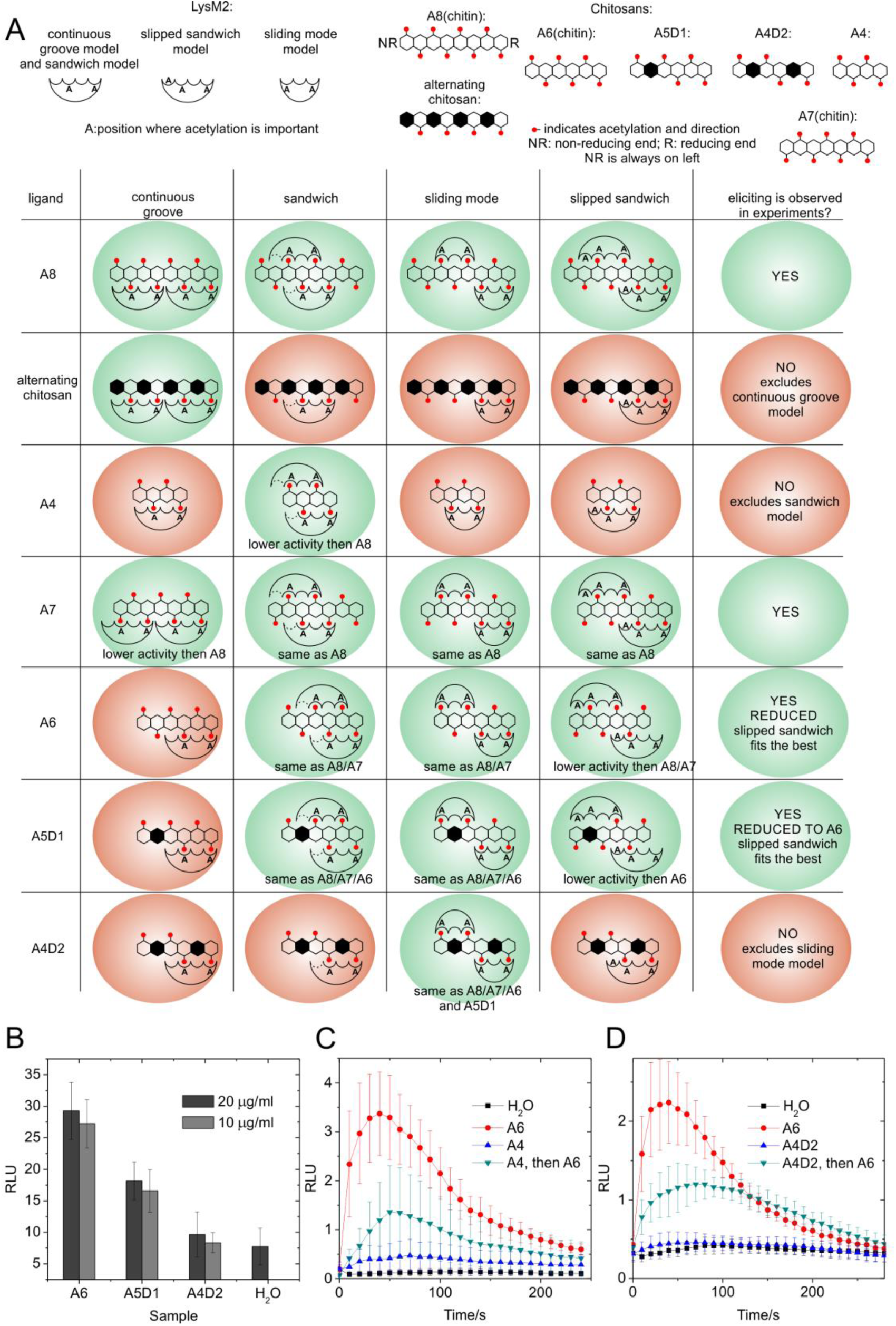
Elicitor activity of chitin oligomers of different DP, and of chitosan oligomers with defined PA. **A**. Expected and observed binding and elicitor activities of chitin/chitosan oligomers to LysM-containing receptors according to the “continuous groove”, “sandwich”, “sliding mode”, and “slipped sandwich” models. The letter “A” in the LysM domain indicates a binding site where acetylation is required for binding. An “A” written inside the domain indicates that the acetyl group must be oriented to face the binding site, one written outside that the acetyl group needs to point away from the protein, as suggested by the 3D structure of the binding pocket in the respective model (see Figure 4 or, for “sliding mode” model, see ^4^). For the “sandwich” model, the weak binding of the first subsite of each LysM ^3^ which is also corroborated by the crystal structure underlying the “sliding mode” model ^4^ is indicated by a dotted line. Predicted productive binding of the oligomers is indicated by green background. Only the “slipped sandwich” model is fully compatible with the existing experimental data. **B**. Eliciting activity of the mono-and double-deacetylated chitosan hexamers ADAAAA and ADAADA; the fully acetylated chitin hexamer served as a positive control. **C**. Inhibiting activity of the fully acetylated chitin tetramer and **D**. inhibiting activity of the double-deacetylated chitosan hexamer ADAADA; the fully acetylated chitin hexamer served as an elicitor. Data given are means ± SD of four replicates from one, representative of three independent experiments.

Similarly, the experimental observation that chitin tetramer acts as an inhibitor ^21^ rather than as an elicitor ^18,22^ argues against the original “sandwich” model, while it is compatible with the other three models (Figure 5A). Then, the experimental evidence shows that chitin heptamer is an equally strong elicitor as chitin octamer, while chitin hexamer is a weak elicitor (Figure 2, ^18,22^). This observation again is not easily compatible with the “continuous groove” model which would instead predict lower activity for the heptamer and no activity for the hexamer. While the “sandwich” model as well as the “sliding mode” model would predict equal activities for these three oligomers, only the “slipped sandwich” model explains the lower elicitor activity of the hexamer. The experimental evidence gained from elicitor assays using fully acetylated chitin oligomers of different DP thus tends to exclude the “continuous groove” and the original “sandwich” model and disfavors the “sliding mode” model, while favoring the “slipped sandwich” model.

To further distinguish between the “sliding mode” and the “slipped sandwich” model, we performed experiments using two defined, partially deacetylated chitosan hexamers. Using a recombinant bacterial chitin deacetylase from *Vibrio cholerae*, VcCDA ^23^, we specifically deacetylated the unit next to the non-reducing end of the chitin hexamer, yielding the mono-deacetylated chitosan hexamer GlcNAc-GlcN-GlcNAc-GlcNAc-GlcNAc-GlcNAc (ADAAAA). An aliquot of this hexamer was then further incubated with a recombinant fungal chitin deacetylase from *Pestalotiopsis spec*., *PesCDA* ^24^ for a short time, yielding the double-deacetylated chitosan hexamer GlcNAc-GlcN-GlcNAc-GlcNAc-GlcN-GlcNAc (ADAADA) as main product with small amounts of ADAAAA and A3D3. The structure and purity of both hexamers were verified using mass spectrometry (Figure E5). When these two chitosan hexamers were tested for elicitor activity, only the mono-deacetylated hexamer was still active, though less than the fully acetylated chitin hexamer, while the double-deacetylated hexamer was elicitor-inactive (Figure 5B). While these experimental results are compatible with the “slipped sandwich” model, they clearly eliminate the “sliding mode” model which would predict full elicitor activity for both of these chitosan hexamers (Figure 5A).

**Figure E5.**
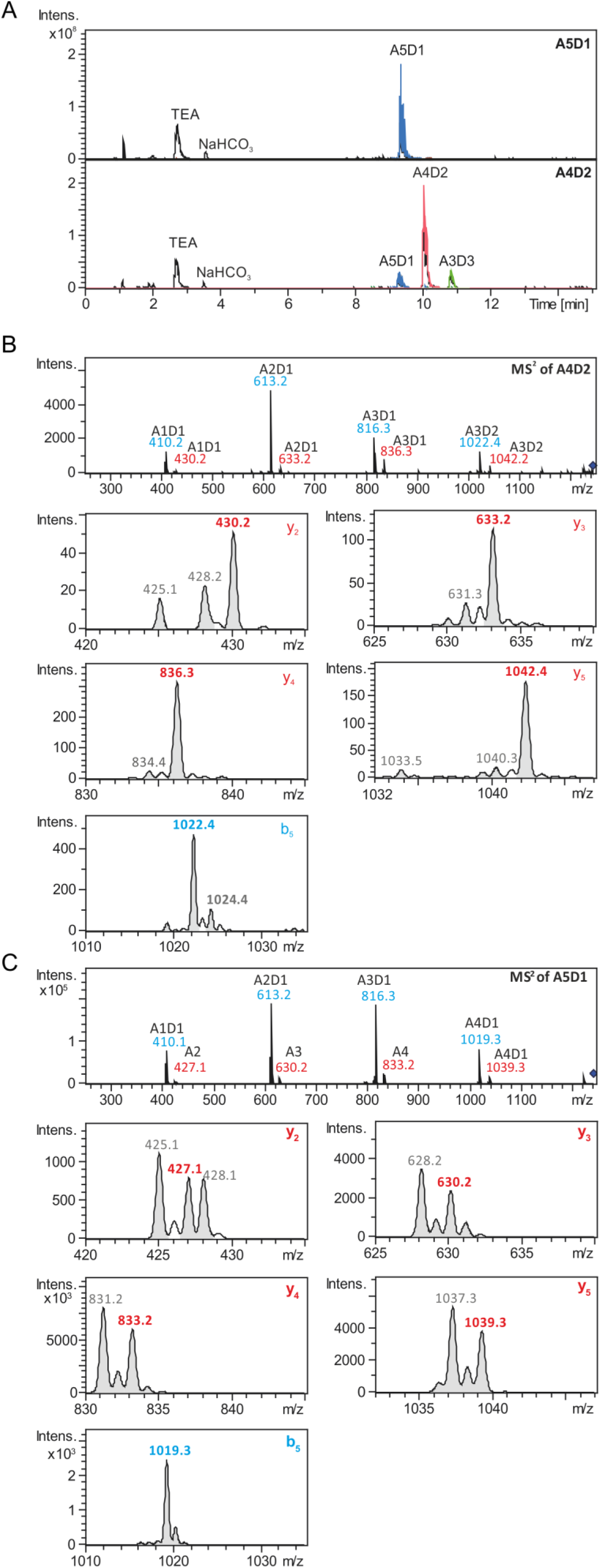
**A**. HILIC-MS base peak chromatogram (black) and extracted ion chromatograms (blue, A5D1; red, A4D2; green, A3D3) of chitosan hexamers produced by incubation of chitin hexamer A6 with (**A5D1**) *Vc*CDA or (**A4D2**) *Vc*CDA followed by *Pes*CDA incubation. The incubation of A6 with *Vc*CDA resulted in the complete conversion of the substrate into A5D1 (> 95%). The sample incubated first with *Vc*CDA followed by *Pes*CDA contained around 75% A4D2, 12.5% A5D1, and 12.5% A3D3. The pattern of acetylation of the **A5D1** and **A4D2** was determined by MS/MS experiments. **B**. Pattern analysis of A4D2 (ADAADA) produced by *Vc*CDA and *Pes*CDA using MS/MS **C**. Pattern analysis of A5D1 (ADAAAA) produced by *Vc*CDA using MS/MS. The samples were [^2^H_3_]*N*-acetylated and ^18^O-labelled to enable quantitative sequencing. The fragments arising from the non-reducing end (b-ions) are shown in blue, fragments arising from the reducing end which contain the ^18^O label (y-ions) are shown in red. The largest b-ion and the y-ions were used to decipher the PA for both chitosan hexamers. The peaks labeled in grey are c-ions.

Finally, we also tested potential inhibitor activities which should be expected for oligomers that can bind to one receptor molecule but are unable to bind a second one in the required conformation to yield an active receptor complex triggering an oxidative burst. For these experiments, we chose the rather weakly elicitor-active chitin hexamer, expecting that potentially antagonistic oligomers will be more readily detectable than when working with a strong elicitor such as the chitin hepta-or octamer. As expected by all four models and as shown previously, the fully acetylated chitin tetramer acted as an inhibitor (Figure 5C). More revealing was the clear inhibitor activity observed for the double-deacetylated chitosan hexamer ADAADA (Figure 5D) confirming that it is bound by AtCERK1 or, possibly, by AtLyK5. As seen in Figure 5B, this hexamer would act as an elicitor in the “sliding mode” model, while acting as an antagonistic inhibitor in the here proposed “slipped sandwich” model, as observed.

Taking all together, unlike (i) the “continuous groove” model which does not explain why the alternating chitosan octamer acts as an inhibitor rather than as an elicitor; unlike (ii) the original “sandwich model” which is at odds with the observation that the chitin tetramer is not an elicitor; and unlike (iii) the “sliding mode” model which does not explain why the double-deacetylated chitosan hexamer ADAADA is not an elicitor but, rather, acts as an inhibitor - our “slipped sandwich” model is in agreement with all the experimentally observed facts.

According to the latest described model of chitin perception in *A. thaliana*, the chitin receptor is formed as a hetero-tetrameric complex of two AtCERK1 and two AtLYK5 proteins (Figure 1B, ^2^). Even though we, like others, were unable to reproduce the reported necessity of AtLYK5 for chitin or chitosan perception (data not shown), we nevertheless tried to accommodate AtLYK5 into our “slipped sandwich” model of the AtCERK1 dimer proposed above. To model the binding groove structure, in contrast to ^2^, we based our model (using SWISS-MODEL workspace ^25^) of AtLYK5 on the available AtCERK1 structure (PDB ID 4EBZ ^1^) rather than on the structure of the fungal chitin binding protein Ecp6 (Figure E1, PDB-ID 4B8V ^9^).

Like AtCERK1 and AtLYK5, Ecp6 possesses three LysM domains, but its structure was found to be significantly different ^9^. In the AtCERK1 structure, three disulfide bonds are formed between different LysM motifs (Figure E6), resulting in a rigid structure, while the four disulfide bonds formed in Ecp6 are intra-rather than inter-motif bonds (Figure E6), allowing much more flexibility and, hence, formation of the ultra-high affinity binding site between LysM1 and LysM3. However, when AtLYK5 is modeled based on the Ecp6 structure ^2,9^, none of these four intra-motif disulfide bonds can be formed. In contrast, when modeling is based on the AtCERK1 structure ^1^, all three inter-motif disulfide bonds can be formed (Figure E6). We therefore conclude that AtLYK5 most likely adopts the same conformation as AtCERK1, OsCEBiP, and, possibly, OsCERK1. In the original model ^2^, chitin binding to the inter-domain binding site buried deep inside one AtLYK5 molecule - or rather the binding of two chitin oligomers to two AtLYK5 molecules - was supposed to “somehow” lead to the binding of two AtCERK1 molecules to the preformed AtLYK5-AtLYK5 homo-dimer, forming the active hetero-tetrameric receptor-ligand complex. Surprisingly, in this original model, chitin binding to AtCERK1 is not even required. However, assuming that AtLYK5 takes the same conformation as AtCERK1, we can easily explain how chitin binding leads to the formation of the AtCERK1-AtLYK5 hetero-dimer and subsequently, due to the preformed AtLYK5-AtLYK5 homo-dimer, to the hetero-tetrameric receptor-ligand complex. Interestingly, and finally, the same dimerization model may then apply for an AtCERK1-AtCERK1 homo-dimer (Figure 6A), an AtCERK1-AtLYK5 heterodimer (Figure 6B), an OsCEBiP-OsCEBiP homo-dimer and, possibly, an OsCEBiP-OsCERK1 hetero-dimer (though no chitin-binding has been shown yet for OsCERK1).

**Figure E6.**
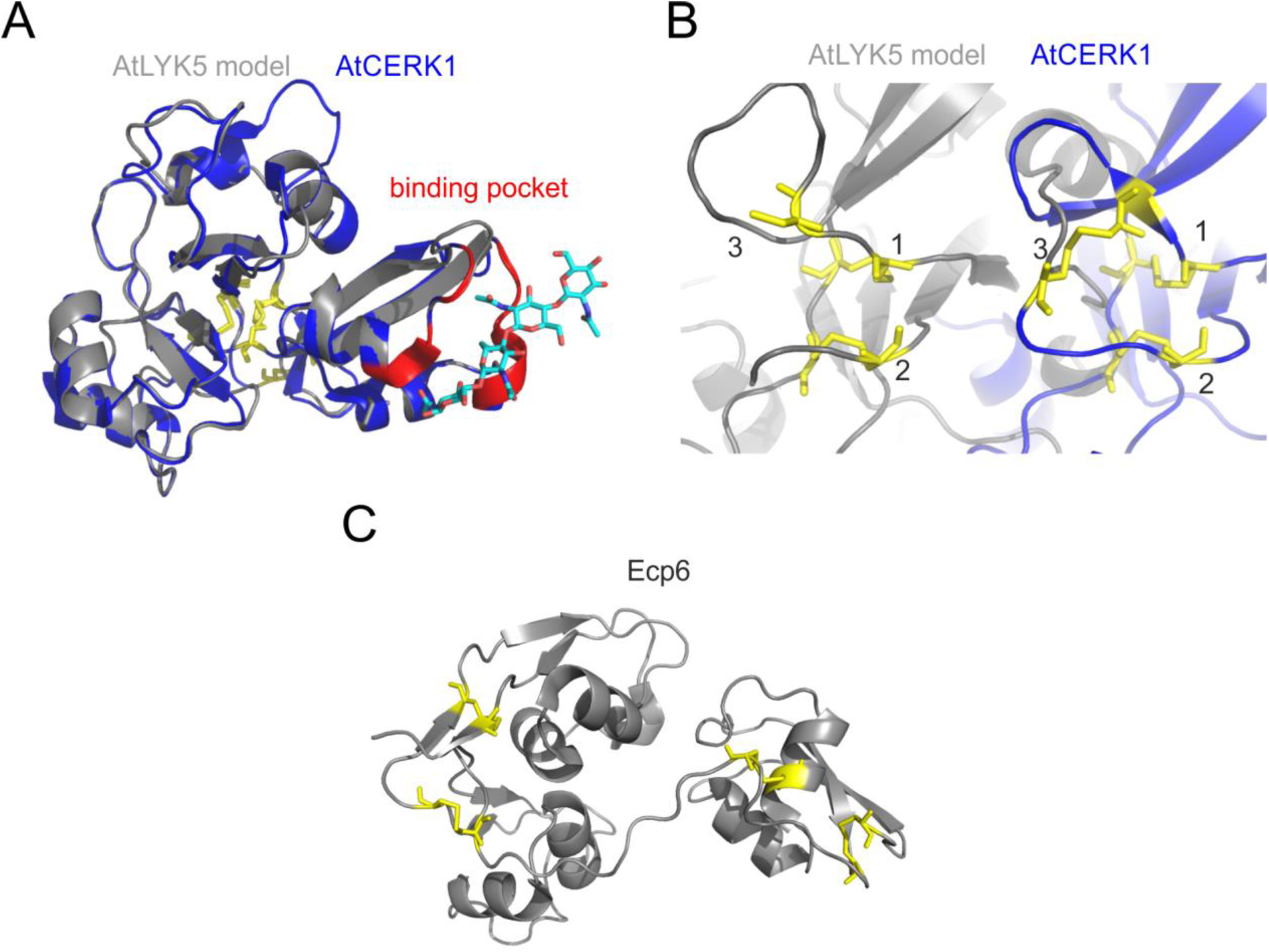
Alternative model for the AtLYK5 ectodomain structure based on the crystal structure of AtCERK1 (PDB ID 4EBZ). **A**. The AtCERK1 structure is given in blue, the AtLYK5 model in grey; the chitin-binding pocket of AtCERK1 is colored in red. **B**. Comparison of the disulfide bonds in the AtCERK1 structure and in the AtLYK5 model. (Bond 3 is not seen in the AtLYK5 model because one of the required cysteines is not modeled, probably because it is the first amino acid visible in the AtCERK1 structure; nevertheless, the bond is expected to be formed in the AtLYK5 protein as well because the corresponding cysteine is present in the sequence at the proper position. **C**. The structure of Ecp6 with its four disulfide bonds is shown for comparison.

**Figure 6.**
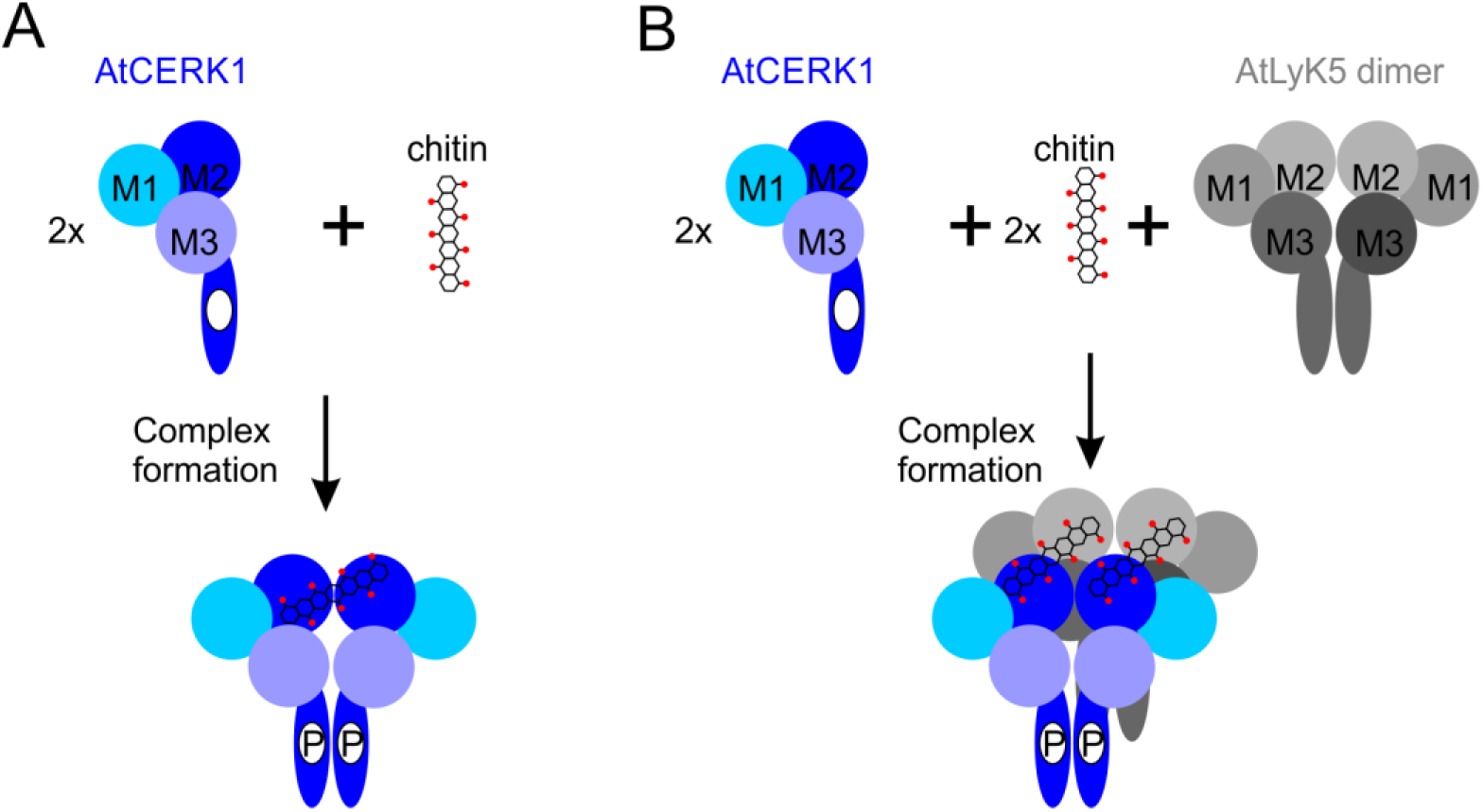
Alternative schemes of receptor complexes in *A. thaliana*. **A**. Two monomers of AtCERK1 bind one molecule of chitin heptamer and form a homo-dimeric receptor complex. **B**. Two monomers of AtCERK1 bind via two chitin heptamers to a preformed AtLYK5 homo-dimer and form a hetero-tetrameric receptor complex. In both cases, dimerization of AtCERK1 within the complex will lead to phosphorylation and downstream signaling.

This improved model for the interaction of LysM-containing chitin-binding proteins and chitin or chitosan oligomers in a “slipped sandwich” mode dimerizing two receptor moieties through their similarly folded LysM2 domains has further implications which we are only briefly mentioning here. The different binding affinities for chitin reported for AtCERK1 at different concentrations ^1,2,21^ suggest that there is a strong cooperativity for chitin binding, indicating that the second AtCERK1 protein would bind much more easily to an AtCERK1-chitin complex than the first AtCERK1 binding to chitin. Similarly, cooperativity can be expected in the AtCERK1-AtLYK5 dimer. As AtLYK5 seems to have a higher affinity for chitin oligomers than AtCERK1 ^2^, this would suggest that the elicitor-active (or inhibiting) ligands would first bind to the existing AtLYK5 dimer which would then easily recruit (or be unable to recruit) two AtCERK1 proteins.

## Acknowledgments

We would like to thank Prof. Naoto Shibuya for initial help with the bioassays and for critically reading the manuscript. This work was funded by DFG in the framework of the Indo-German International Research Training Group on Molecular and Cellular Glyco-Sciences - MCGS, through doctoral fellowships to EG and RM.

## Materials and Methods

### Seedling preparation

Arabidopsis seeds of the wildtype Col-0 and its AtCERK1 knock-out mutant *cerk1-2* ^20^ were sterilized with 1 ml of 70% ethanol for 1 min, then ethanol was substituted with 20% sodium hypochlorite solution containing 1% Tween 20. Seeds were washed 4-5 times with sterile water and plated aseptically on 0.1% agar, containing 2.2 g L^-1^ Murashige and Scoog (MS) salts (Deutsche Phytoengineering Chemical Company, Langenberg, Germany) supplemented with 1% sucrose, at pH 5.7. Seedlings were grown in Percival AR-66 L4 growth chambers (Percival Scientific, Perry, USA) at 22°C in an 8-h light/16h-dark photoperiod.

### Luminol chemiluminescense assay for H_2_O_2_ detection in 96 well plates

To monitor the H_2_O_2_ accumulation, we modified the method of ROS measurement after ^26^ that quantifies luminol chemiluminescence as described previously ^27^. Seven-day old seedlings were transferred individually into single wells of a 96-well microtitre plate (Thermo Fisher Scientific Nunc A/S, Denmark), containing 170 μl of nutrients solution ^28^ complemented with 1% sucrose per well. Using a multichannel pipette, 30 μl of Luminol L-012 (Waco Pure Chemical Industries, Neuss, Germany) at the stock concentration of 100 μg/ml was quickly added to each well. Then, the plate was incubated for 3 h at 22°C, covered with aluminium foil to protect the luminol from degradation. Directly before the measurement, elicitors were added to each well and the plate was placed in the microtitre plate reader (Luminoskan™ Ascent Microplate Luminometer, Thermo Scientific, Schwerte, Germany). For the detection of reactive oxygen species released by *Arabidopsis* seedlings, every well was measured for 5 s every 10 min for 10 h. Kinetics of H_2_O_2_ production was determined by integration of data for every well over the reading period. Every time point is the mean value of four seedlings from four wells (either mock or chitosan elicited). All experiments were repeated independently at least three times.

To assess possible inhibitory activities of chitin/chitosan oligomers, seedlings were preincubated with potential antagonists (60 μM) for 15 min in the microtiter plate, followed by addition of chitin hexamer as an elicitor (10 μM), as described above.

### Preparation and characterization of chitin and chitosan oligomers

Purified chitin (DA 100%) and chitosan (DA 0%) oligomers with DP ranging from 3 to 8 were prepared by HCl hydrolysis of α-chitin (Mahtani Chitosan, Veraval, India; Batch 25) or DA 0% chitosan (Mahtani Chitosan, Veraval, India; batch 134), respectively, as described ^29^. Separation of oligomers was achieved using size exclusion chromatography (SEC), as described ^30^. Purified oligomers were analyzed using UHPLC-ELSD-ESI-MS ^31^.

### Preparation and characterization of partially N-acetylated chitosan polymers

Polymeric chitosans (average DP 1300) with DAs ranging from 1% to 60%, in which the GlcNAc and GlcN residues are randomly distributed along the polymer chain, were prepared and analysed as described before ^32^.

### Preparation and characterization of partially N-acetylated chitosan oligomers

Purified partially acetylated chitosan oligomers were produced by enzymatic hydrolysis of partially re-N-acetylated chitosan with DA 10% and 35% as described above. Purified recombinant *Bacillus licheniformis* chitinase ^33^ and *Bacillus sp*. chitosanase ^34^ were used to produce chitosan oligomers with DA 10% and DA 35%, respectively (Sven Basa et al., unpublished). Oligomers were separated using SEC ^32^ and analyzed using MALDI-TOF MS ^35^.

Defined mono-and double-deacetylated chitosan hexamers were produced by enzymatic deacetylation of fully acetylated chitin hexamer (Megazyme, USA). Purified recombinant chitin deacetylases *Vc*CDA from *Vibrio cholerae* ^23^, and *Pes*CDA from *Pestalotiopsis spec*. ^24^ were used to produce GlcNAc-GlcN-GlcNAc-GlcNAc-GlcNAc-GlcNAc (*Vc*CDA only) or GlcNAc-GlcN-GlcNAc-GlcNAc-GlcN-GlcNAc (first *Vc*CDA followed by *Pes*CDA incubation) chitosan hexamers. The reactions were stopped by removing the enzymes with a 3K-Pes-filter (VWR, Darmstadt, Germany) at 4°C, centrifuging at 13000 rpm for 30 min. The target hexamers were analysed using HILIC-ESI-MS ^31^.

